# Cell volume regulates terminal differentiation of cultured human epidermal keratinocytes

**DOI:** 10.1101/2024.12.09.627463

**Authors:** Sebastiaan Zijl, Toru Hiratsuka, Atefeh Mobasseri, Mirsana Ebrahimkutty, Mandy Börmel, Sergi Garcia-Manyes, Fiona M. Watt

## Abstract

Differentiation of cultured human epidermal stem cells is regulated by interactions with the underlying substrate. Whereas differentiation is typically stimulated when keratinocytes are prevented from spreading, we previously identified two micron-scale topographical substrates that regulate differentiation of spread cells. On one substrate (S1), individual cells interact with small circular topographies, and differentiation is stimulated; on the other (S2), cells interact with larger triangular topographies, and differentiation is inhibited. By scanning electron microscopy we visualised substrate interactions at higher resolution than previously and using live cell imaging we established that induction of the differentiation marker involucrin did not involve transient cell rounding on S1. Bulk gene expression profiling did not reveal any differences between cells on S1 and S2 prior to the selective upregulation of differentiation markers at 12h on S1 and cell stiffness was lower on both S1 and S2 than on flat substrates. Nevertheless, cells on S2 differed from cells on flat and S1 substrates because they exhibited reduced cell volume, prompting us to explore whether cell volume could regulate differentiation independent of culture substrate. Treatment with polyethylene glycol (PEG) reduced cell volume and inhibited differentiation regardless of whether keratinocytes were seeded on flat, S1 or S2 substrates, micropatterned islands or in suspension. Conversely, treatment with deionised water increased cell volume and stimulated differentiation of substrate adherent keratinocytes. On flat substrates treatment with the Ca^2+^ chelator 1,2-bis-(2-aminophenoxy)ethane-N,N,N’,N’-tetraacetic acid acetoxymethyl ester or an inhibitor of the water channel aquaporin 3 blocked induction of differentiaton by deionised water, whereas the gadolinium^3+^, a stretch-activated calcium channel blocker, did not. Our studies identify a new mechanism by which keratinocyte-niche interactions regulate initiation of differentiation.

## Introduction

Human skin comprises the epidermis, which is formed of multiple layers of epithelial cells called keratinocytes, and the underlying connective tissue, or dermis (Rognoni and Watt, 2018). The epidermis and dermis are separated by a basement membrane. The basal epidermal layer, which is attached to the basement membrane, contains a patterned distribution of stem cells and cells that have initiated terminal differentiation. Differentiating cells detach from the basement membrane and move through the suprabasal epidermal layers to the skin surface, from which they are shed (Zijl et al., 2022).

Human epidermal keratinocytes can be grown in culture under conditions that support maintenance of stem cells and the process of terminal differentiation (Zijl et al., 2022). Cultured human epidermis can therefore be used to examine, at single cell resolution, how stem cells make fate decisions based on signalling cues from the external microenvironment (Watt, 2016; Louis et al., 2022; Negri and Watt, 2022; Negri et al., 2023).

Previous studies have highlighted the role of cell-substrate interactions in controlling differentiation of human epidermal stem cells (Zijl et al., 2022). When single cells are seeded on ECM-coated micro-patterned islands, differentiation is triggered by restricted spreading, which is dependent on the ratio of F- to G-actin and activation of serum response factor (SRF) (Watt et al., 1988; Connelly et al., 2010). Differentiation is also triggered when cells are plated on ECM coated soft hydrogels or hydrogel-nanoparticle composites with high nanoparticle spacing (Trappmann et al., 2012). On soft hydrogels differentiation is mediated by downregulation of extracellular signal-regulated kinase (ERK)/mitogen-activated protein kinase (MAPK) activity.

Whereas keratinocyte differentiation is typically associated with reduced cell spreading, we have also found that micron-scale substrate topographies can promote differentiation of spread cells (Zijl et al., 2019). In particular we have fabricated a substrate (designated S1), comprising circular topographical features of approximately 3 µm diameter, 5 µm height and unequal spacing, that promotes differentiation of spread cells via a mechanism that is blocked by Rho kinase inhibition or treatment with the myosin II inhibitor blebbistatin, but not by SRF inhibition. Conversely, a substrate comprising regularly spaced right-angled triangles (8 µm sides) of 5 µm height (designated S2) suppressed differentiation relative to flat surfaces. In the present study we have explored how keratinocytes respond to each of these substrates, uncovering a potential role for cell volume in regulating keratinocyte differentiation.

## Results

### Keratinocytes on S1 substrates can initiate differentiation while spread

In order to enrich for undifferentiated keratinocytes (stem cells) we seeded single cell suspensions, comprising a mixture of basal and differentiated cells, on collagen-coated S1, S2 or flat surfaces for 1 h and washed off the non-adherent cells (Jones and Watt, 1993; Zijl et al., 2019). Scanning electron microscope (SEM) images of keratinocytes 24h after plating (Figure 1) revealed that cells on the S1 substrates spread over the individual pillars (Figure 1C, D); some pillars at the edges of the substrates were bent, suggesting that they had been subject to pulling force by the cells (Figure 1D). The cytoplasm of cells on S2 appeared to be draped over the triangular features (Figure 1E, F). In some cells on the S2 substrates, the nucleus extended above the surface of the triangular features (Figure 1F), while in other cells the nucleus appeared to have been accommodated between the triangular features (Figure 1E), consistent with the nuclear distortion observed by light microscopy (Zijl et al., 2019). Heterogeneity in terms of flattened versus protruding nuclei was also evident on flat and S1 substrates (Figure 1A-D).

**Figure 1.**
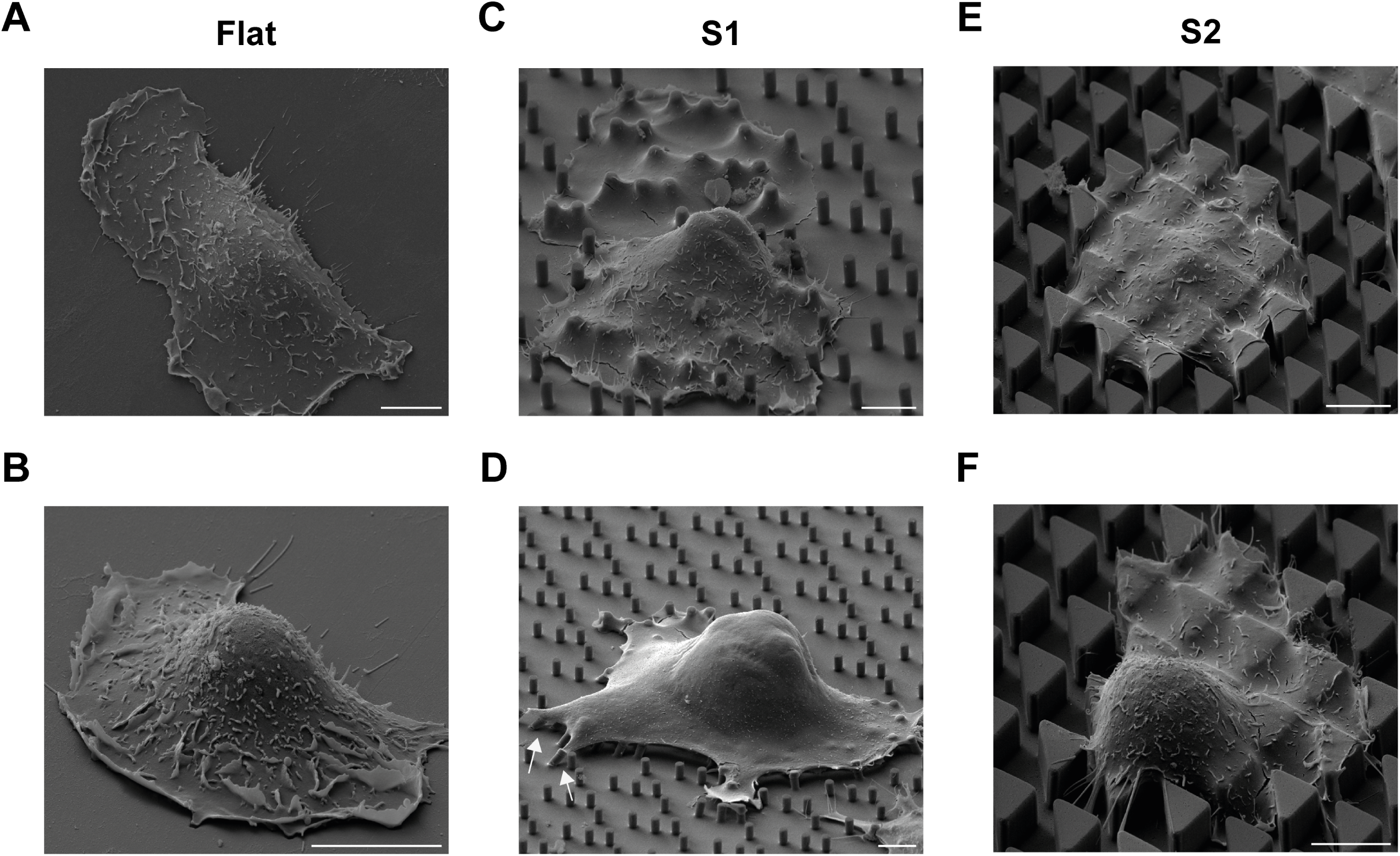
SEM of single keratinocytes on flat, S1 and S2 substrates. Single spread cells on flat (A, B), S1 (C, D) and S2 (E, F) substrates. Some S1 substrate pegs at the cell periphery were bent (arrows, D). In some cells on S2 the nucleus was visible above the substrate surface (F), whereas in others (E) it was not. Scale bars: 10 μm.

To confirm that cells seeded on S1 initiated differentiation without rounding up we performed live imaging of keratinocytes transduced with a pLenti-IVL-mCherry-LifeAct-EGFP reporter. The reporter expresses LifeAct-EGFP, which binds to actin filaments (Riedl et al., 2008; Belin et al., 2014), under the control of the hPGK promoter, and mCherry under the control of the involucrin promoter (Hiratsuka et al., 2020) (Figure 2A). Flow cytometry confirmed that mCherry was expressed by cells of high forward and side scatter (enriched for differentiating cells; Jones and Watt, 1993) and that mCherry-positive cells co-expressed GFP (Figure 2A). We flow sorted IVL-LifeActGFP+ cells (Figure 2A) and seeded them on S1 (Figure 2B).

**Figure 2.**
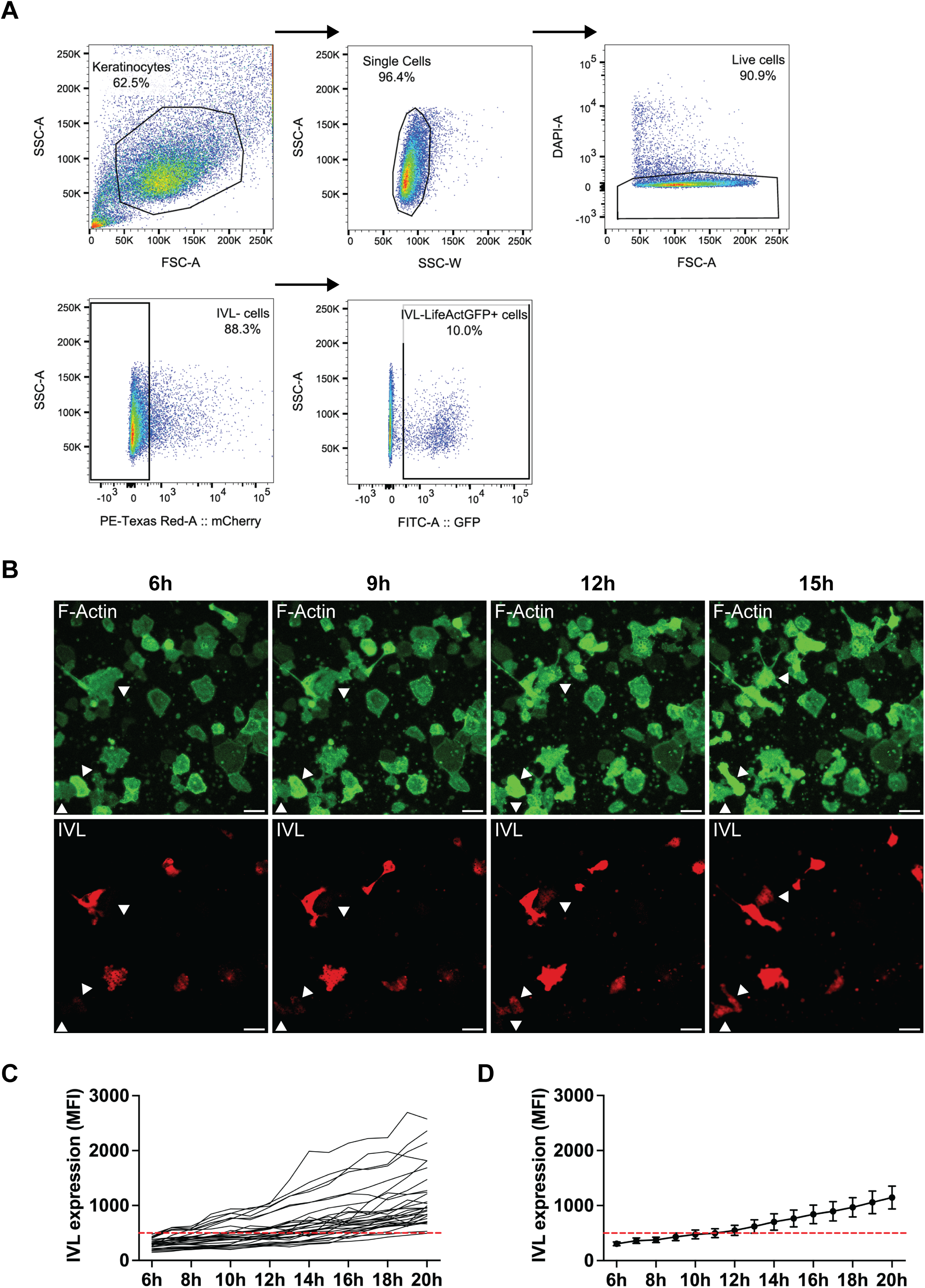
Expression of INV-mCherry reporter by keratinocytes seeded on S1 culture substrates. A. Overview of the FACS sorting strategy to select INV-mCherry negative cells expressing GFP (IVL-LifeAct-GFP+ cells) prior to plating on the substrates. Each plot represents a sequential gating step. Arrows indicate the gating sequence. Keratinocytes were first gated by forward (FSC-A) and side scatter (SSC-A), then for single cells and then live cells (DAPI-negative). mCherry-negative (IVL-negative) cells were selected for seeding. In the final gate the co-expression of mCherry and GFP by cells with high SCC is shown. (B-D) IVL-LifeAct-GFP+mCherry-cells selected as in (A) were plated on S1 substrates for 45-60 minutes and non-adherent cells were washed off. Adherent cells were monitored by live cell imaging. (B) Representative field of cells. Two cells marked with arrowheads initiated mCherry expression while spread. (C, D) 30 cells that initiated mCherry expression during the imaging period were pooled from three independent experiments for analysis. The IVL-mCherry intensity of each cell was measured by calculating the mean fluorescence intensity (MFI) of cells at different time points. Cells with an MFI > 500 were considered to be undergoing differentiation. (C) Traces of individual cells are shown. (D) Mean mCherry signal of all 30 cells. Error bars: 95% confidence interval (CI). Dashed line shows mean fluorescence of all 30 cells throughout the time course. Scale bars: 50 μm.

Cells expressing IVL-mCherry-LifeAct-EGFP were imaged at 1h intervals starting 4-6h after seeding. Figure 2B shows representative images of a single field of keratinocytes. At 6h several cells in the field were already expressing mCherry and exhibited variable morphologies. Two cells (arrowheads) were mCherry-negative at 6h and 9h but expressed mCherry at 12h and 15h; in each case the onset of mCherry expression occurred while the cells were spread.

Using LifeAct-EGFP to visualise individual cells, whether or not they had differentiated, we measured the mean mCherry fluorescence of 30 cells on S1 that had upregulated mCherry by 20h (Figure 2C). The mean mCherry signal from cells increased above baseline from 12h onwards (Figure 2D), consistent with earlier studies of the kinetics of keratinocyte differentiation in culture (Connelly et al., 2010; Zijl et al., 2019).

These observations establish that cells plated on S1 can upregulate involucrin expression while spread.

### RNAseq does not reveal differences in overall gene expression between cells on S1 and S2 prior to differentiation

Changes in gene expression associated with commitment to differentiation have previously been described in keratinocytes undergoing differentiation in single cell suspension (Mishra et al., 2017), including a network of interacting protein phosphatases that are also upregulated during the basal to suprabasal transition of keratinocytes in healthy human skin (Reynolds et al., 2021; Negri et al., 2023). To examine whether there were unique features of the gene expression programme of keratinocytes undergoing commitment on S1, we seeded keratinocytes on S1 or S2 substrates and extracted RNA at 1h (initial adhesion), 4h (differentiation commitment) and 12h (increased differentiation on S1 versus S2). RNA was harvested in three independent experiments and subject to bulk RNA sequencing (RNAseq) (Figure 3A). In total 18 samples were submitted for RNA seq (3 time points in 3 independent experiments on each substrate). Sequencing and bioinformatic analysis were performed by Genewiz, Inc.

**Figure 3.**
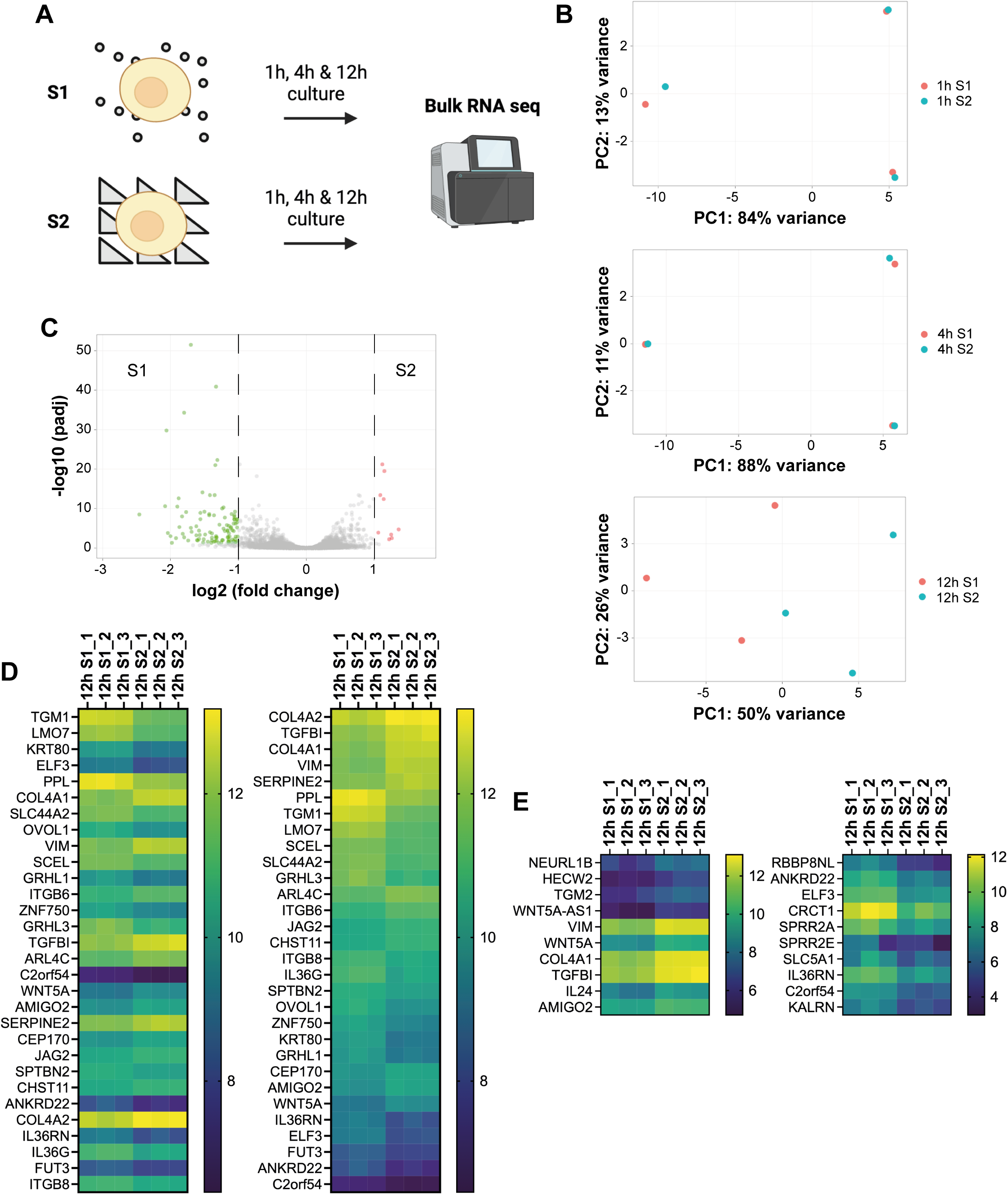
Bulk RNA sequencing of keratinocytes seeded on S1 or S2 substrates. (A) Schematic overview. Cells were cultured for 1h, 4h or 12h in complete FAD medium prior to RNA extraction. RNA was extracted from cells in three independent experiments (replicate experiments are denoted e.g. 1h S1_1, 1h S1_2 and 1h S1_3). Schematic was created using BioRender.com. (B) Principal component analysis (PCA) plots comparing cells on S1 (red dots) with cells on S2 (blue dots) at different time points. Each data point in the graph represents a different sample. PC: principal component. (C) Volcano plot showing global transcriptional differences between cells on S1 and S2 at 12h. Each data point represents one gene. The log2 fold change (LFC) of each gene is shown on the x-axis and the -log10 of its Benjamini-Hochberg adjusted p-value (padj) is shown on the y-axis. Genes with a padj < 0.05 and LFC > 1 (red dots) are upregulated on S2 Genes with padj < 0.05 and LFC < -1 (green dots) are downregulated on S2. (D, E) The most significantly differentially expressed genes at 12h. (D) Heatmaps of the 30 most significantly differentially expressed genes (DEGs) on S1 and S2 at 12h. Genes were sorted by their Benjamini-Hochberg adjusted p-value (padj) (left hand column) or expression based on log2 transformed expression values and log2 fold change in expression (right hand column). Left hand column: the most significantly DEGs (lowest padj) are shown first. Right hand column: highly expressed genes upregulated on S2 (compared to S1) are shown first, followed by highly expressed genes downregulated on S2. Each square represents the (log2 transformed) expression level for a different gene (rows) or sample (columns). (E) Heatmaps for the top 10 upregulated genes (highest LFC) (left hand column) and downregulated genes (highest negative LFC) (right hand column) on S2 at 12h (compared to S1). Genes were selected based on a padj of < 0.05 and ranked according to their log2 fold change (LFC). Each square represents the log2 transformed expression level for a different gene (rows) or sample (columns).

Principal component analysis (PCA) revealed that cells on S1 and S2 had highly similar gene expression at 1h and 4h, but diverged at 12h (Figure 3B). At 1h and 4h the differences between cells on S1 and S2 were lower than the differences between different experiments (technical replicates) (Figure 3B). Nevertheless, cells on S1 and S2 showed significant differences in gene expression at 12h (Figure 3B). The majority of differentially expressed genes (100/109 genes, 92%) at 12h were downregulated on S2 (log2 fold change > 1or < -1, padj < 0.05) (Figure 3C and Supplementary Table 1). This was also the case when we lowered the log2 fold change (LFC) threshold to LFC > 0.5/LFC < -0.5 (242 out of 337 genes, 72%; Supplementary Table 1). The full list of differentially expressed genes (LFC > 0.5 or LFC < -0.5, padj < 0.05) can be found in Supplementary Table 1.

As shown in Supplementary Figure 1, the top GO terms for genes that were differentially expressed between S1 and S2 at 12h were ‘epidermis development’, ‘keratinocyte differentiation’, ‘establishment of skin barrier’ and ‘peptide crosslinking’ (which includes genes such as TGM1 that are involved in forming the epidermal barrier (Steinert and Marekov, 1999; Candi et al., 2005) and keratinization (a process also linked to differentiation; Redmond and Coulombe, 2021) (Figure 3D). Thus the majority of the genes corresponded to genes that are upregulated during keratinocyte terminal differentiation, consistent with our earlier findings (Zijl et al., 2019). The top three genes upregulated on S2 at 12h were TGFBI, COL4A1 and WNT5A, all of which are known to be expressed in the basal epidermal layer (Romanowska et al., 2009; Li et al., 2022) (Figure 3E). VIM is expressed at low levels by cultured keratinocytes (Kariniemi et al., 1982) and was upregulated on S2 compared with S1. Real-time quantitative PCR of differentially expressed genes at 12h confirmed that differentiation was upregulated on S1 and downregulated on S2 compared to the flat substrates, while WNT5A, TGFBI and VIM were selectively upregulated on S2 (Supplementary Figure 2A, B).

We conclude that using bulk RNAseq we could confirm the stimulation of differentiation on S1 but could not identify a distinct commitment programme.

### Effects of S1 and S2 on cell volume and stiffness

We next compared cell and nuclear volume on the different topographies by confocal microscopy of keratinocytes labelled with anti-keratin 14 (K14) and DAPI (Guo et al., 2017; Hansen et al., 2022; Koushki et al., 2023; Li et al., 2021) (Figure 4A). Cells on S2 had a significantly lower total volume at both 4h and 12h (Figure 4B, C). At 4h, the nuclear volume was also lower on S2 (Figure 4D) although this was not the case at 12h (Figure 4E). Since the cell volume on S1 was similar to that on flat substrates (Figure 4B, D), the changes in cell volume were specific to S2.

**Figure 4.**
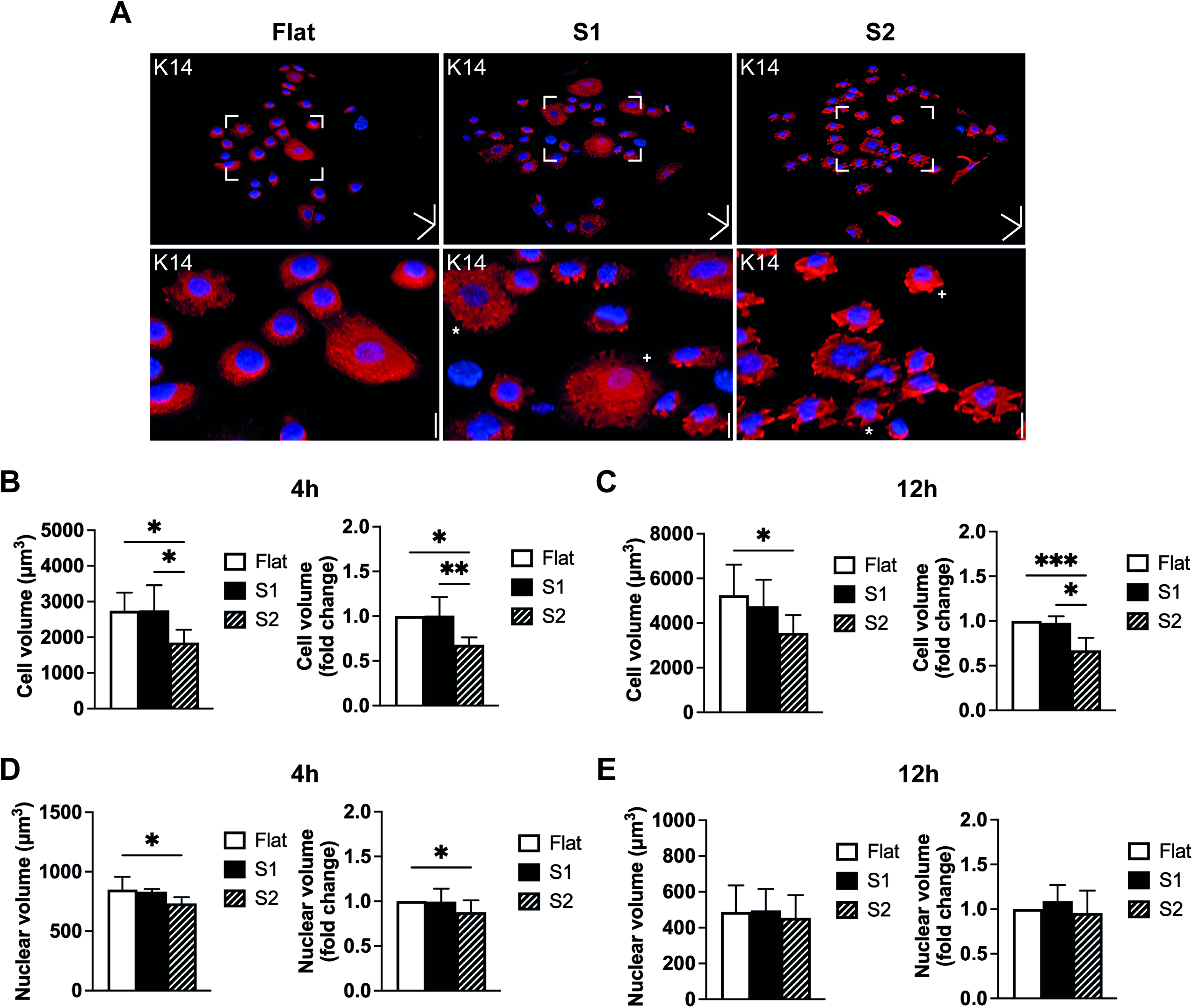
Volume of cells seeded on different substrates. (A) Representative 3D reconstruction images of anti-keratin-14 (K14, red) and DAPI (blue) labelled cells grown on Flat, S1 and S2 substrates for 4h. Boxes: areas of interest shown at higher magnification below. Scale bars: 50 μm (x, y and z direction), 20 μm (zoom-in images, z direction). Cell (B, C) and nuclear (D, E) volume measurements on flat, S1 and S2 substrates after 4h (B, D) or 12h (C, E) of culture. Total and fold change (normalised to flat substrates) in volume are shown. Data are from three independent experiments in which >300 cells were analysed per experiment. Three technical replicates were measured per substrate. Statistics: one-way ANOVA (+ Tukey’s) test (absolute volumes), Kruskal-Wallis (+ Dunn’s) test (fold changes). Error bars: mean + SD. * p-value < 0.05, ** p-value < 0.01, *** p-value < 0.001.

It has previously been reported that changes in cell volume affect cell stiffness (Guo et al., 2017). In addition, actomyosin regulates differentiation on S1 (Zijl et al., 2019) and an increase in cortical tension can increase cell stiffness (Cartagena-Rivera et al., 2016; Harris et al., 2012). We therefore performed atomic force microscopy (AFM) on individual keratinocytes (Figure 5A) to determine whether the differential effects of S1 and S2 on differentiation were correlated with differences in cell stiffness. These experiments showed that cells on S1 and S2 were significantly softer than cells on flat controls (Figure 5B-E), ruling out an association between cell stiffness and differentiation on S1 and S2.

**Figure 5.**
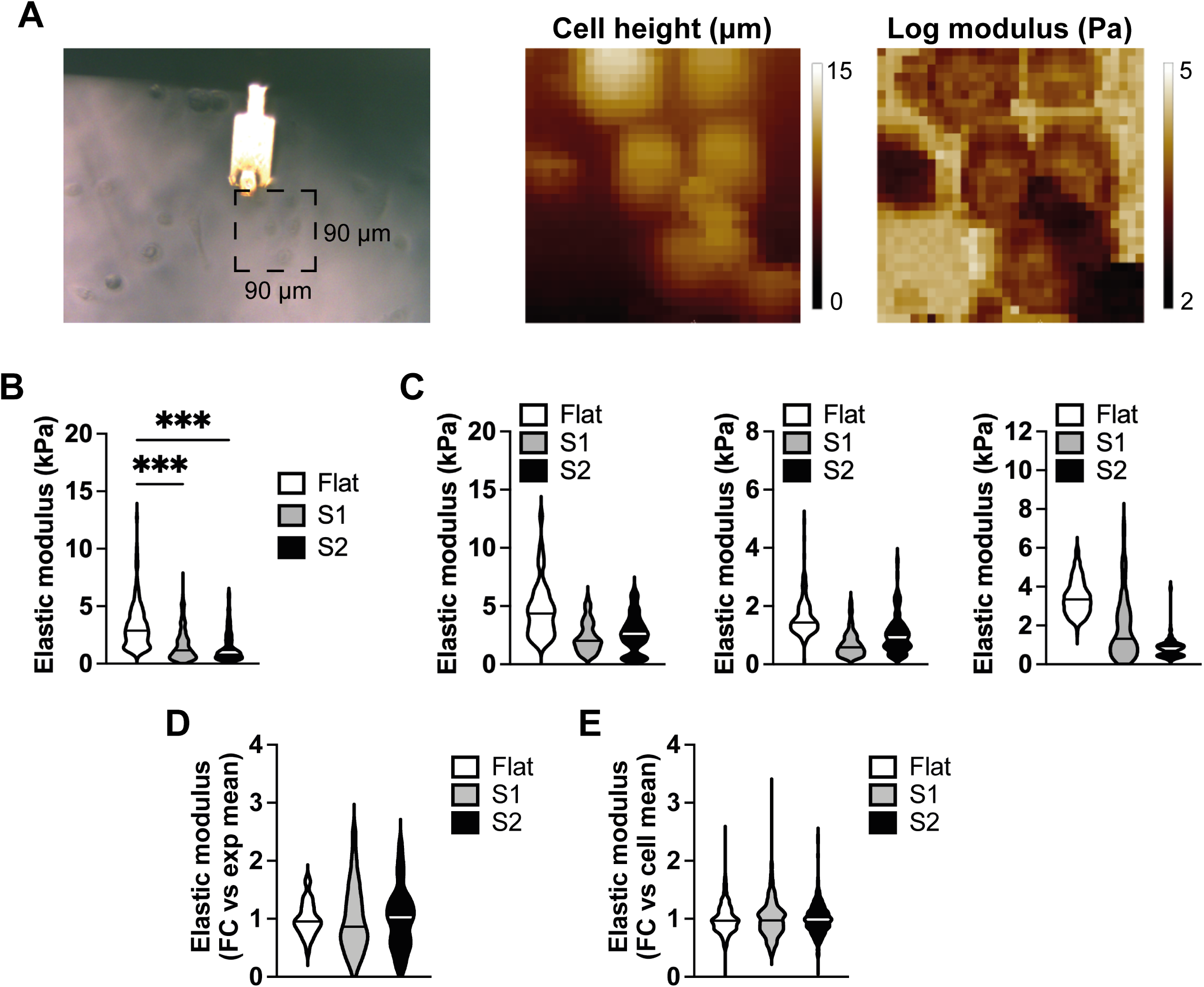
Stiffness measurements of cells seeded on different substrates. (A) Representative brightfield image (left hand side) of an AFM probe measuring bulk elastic modulus (cell stiffness) of cells on a flat substrate. Height (middle panel) and log modulus (right hand side) measurements are shown. (B) Quantification of cell stiffness of cells grown on flat, S1 and S2 substrates for 12h (*n* = 3 independent experiments). 10 randomly chosen cells were analysed per experiment (20 measurements per cell). (C) Data from the individual experiments shown in (B). (D) Variation in cell stiffness within individual experiments shown in (C). Values normalised to the mean (1.0) of individual experiments. (E) Variation in stiffness between individual cells shown in (C). Values normalised to the mean (1.0) of individual cells. Plots (B-E) represent the median (horizonal line) and range in values. FC: fold change; kPa: kilopascal. *** p-value < 0.001.

### PEG and DI regulate keratinocyte differentiation on flat surfaces and in suspension

The observation that cells exhibited a reduced volume on S2 substrates that inhibit differentiation led us to investigate whether changes in cell volume could directly influence differentiation. We examined the effects of adding the molecular crowding agent polyethylene glycol 300 (PEG 300), which decreases cell volume, or deionized water (DI), which increases cell volume (Cai et al., 2019; Guo et al., 2017; Li et al., 2021; Rashid et al., 2023; Tomba et al., 2022). Cells were treated with PBS as a control. Cells were allowed to adhere to a flat substrate for 1h then treated with PEG 300, DI or PBS for 3h prior to fixation and labelling (Figure 6). These experiments confirmed that PEG 300 reduced and DI increased cell volume relative to PBS (Figure 6B). PEG 300 treatment also led to a small reduction in nuclear volume (Figure 6C).

**Figure 6.**
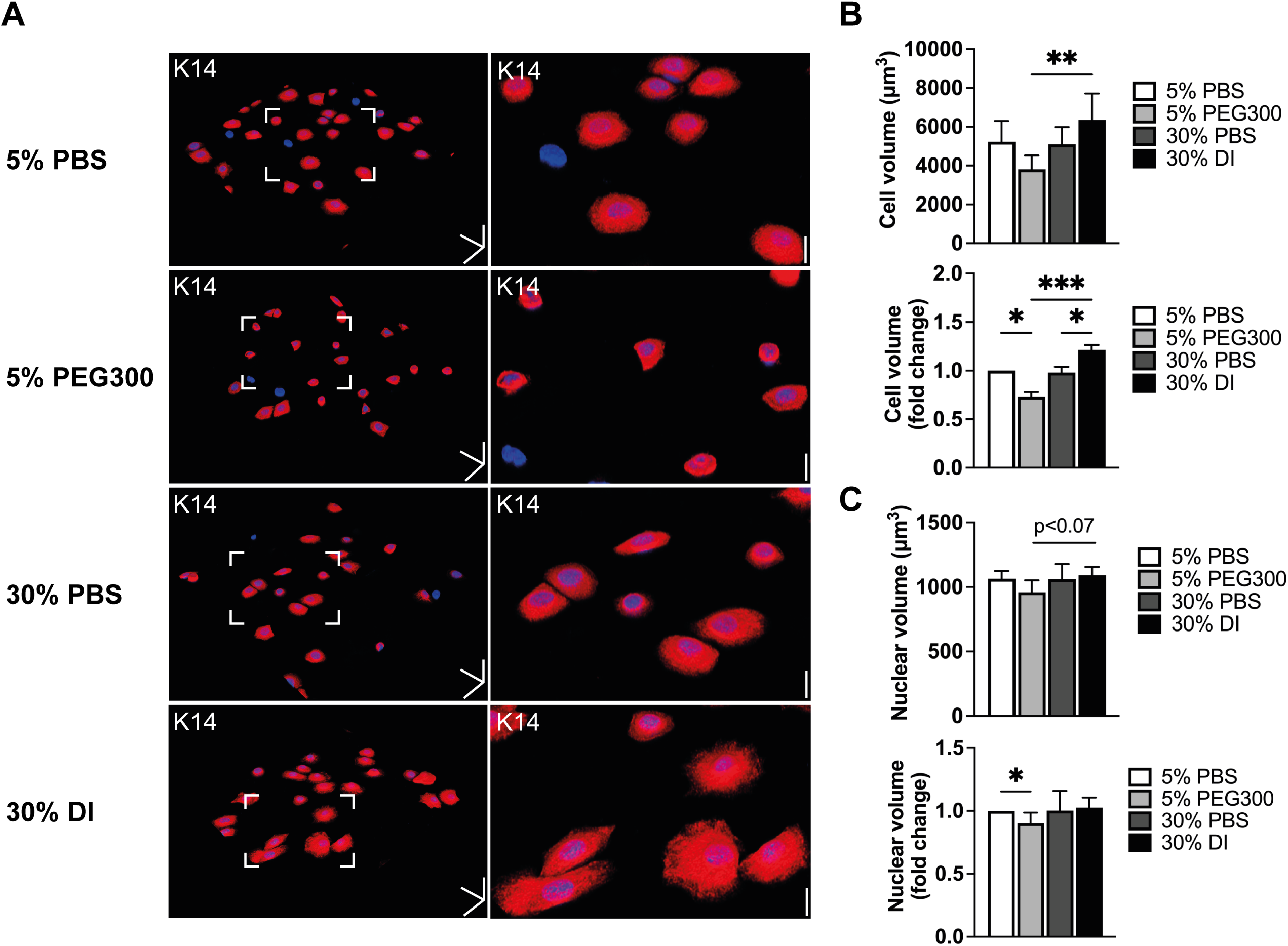
Modulation of cell volume on flat substrates. (A) Representative 3D reconstruction images of cells labelled with DAPI (blue) and anti-keratin14 (K14) after treatment with 5% PBS, 5% PEG 300, 30% PBS and 30% DI for 3h. Boxed areas of interest in left hand panels are enlarged in right hand panels. Scale bars: 50 μm (x, y and z direction), 20 μm (zoom-in images, z direction). (B) Quantification of cell volume and fold change in cell volume (normalised to 5% PBS). (C) Quantification of nuclear volume and fold change in nuclear volume (normalised to 5% PBS). Data from three independent experiments in which >300 cells were analysed per experiment. In each experiment three technical replicates were analysed per condition. Statistics: one-way ANOVA (Šídák’s) test (volume measurements), Kruskal-Wallis (+ Dunn’s) test (fold changes). Error bars: mean + SD. PBS: phosphate-buffered saline, PEG300: polyethylene glycol 300, DI: deionised water. * p-value < 0.05, ** p-value < 0.01. *** p-value < 0.001.

24h treatment with PEG reduced differentiation on flat substrates, while DI stimulated differentiation (Figure 7A-C). Treatment with PEG 300 caused a striking change in keratinocyte morphology (Figure 7B). There was a stronger effect of 5% compared to 2% PEG 300 (Figure 7D). PEG 300 also inhibited differentiation of cells in suspension, as demonstrated by Western blotting (Fig. 7E; Supplementary Figure S3). The same effects on differentiation were observed on micropatterned islands, under conditions in which spread area – determined by the micropatterns – did not change (Figure 8).

**Figure 7.**
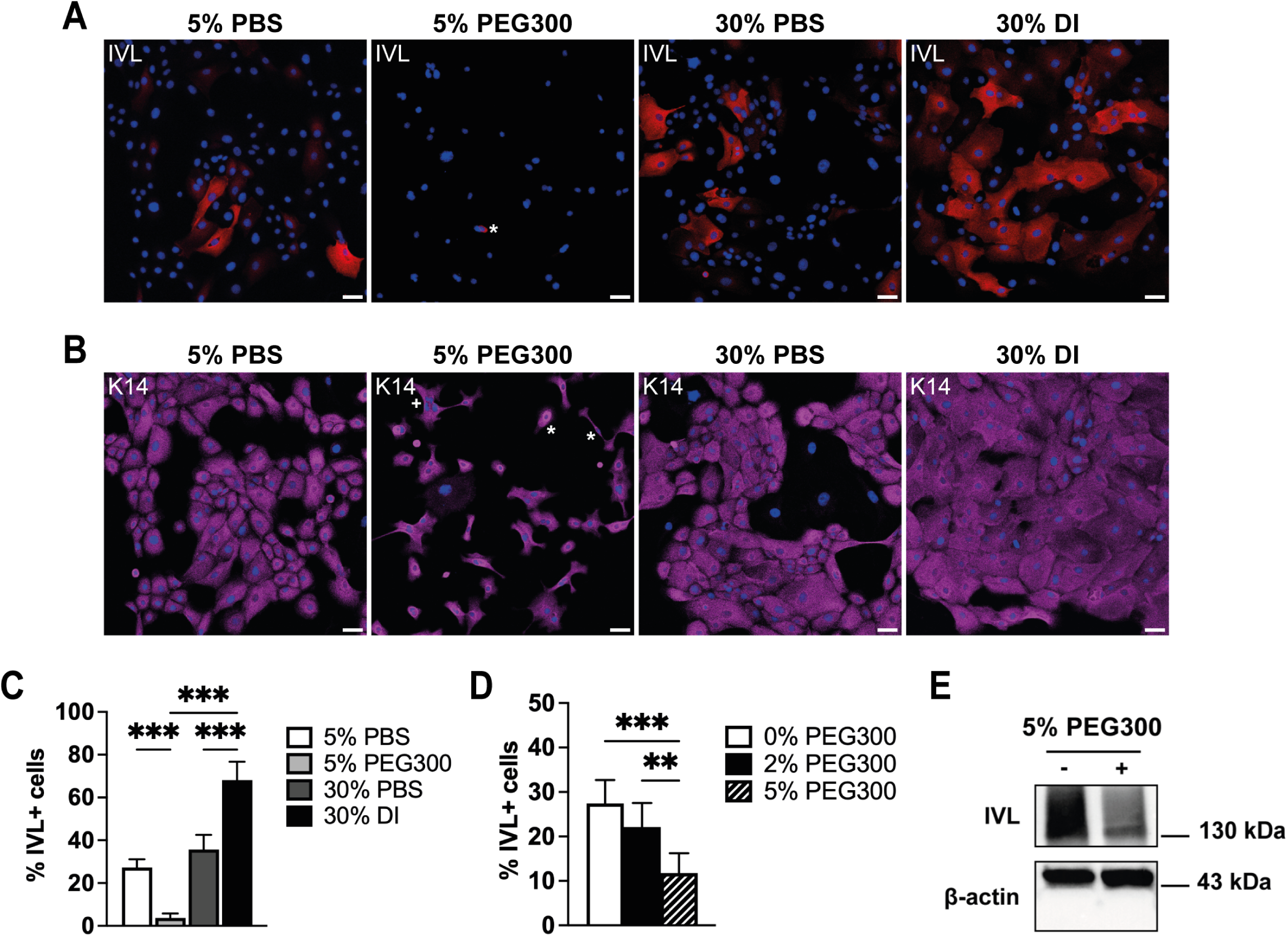
Modulation of differentiation of adherent and suspended keratinocytes with PEG 300 and DI. (A, B) Representative images of anti-involucrin (A: IVL, red) and anti-keratin 14 (B: magenta) labelled cells with DAPI counterstain (blue). Cells were grown for 24h on flat substrates (unpatterned 96-well plates) in the presence of 5% PBS, 5% PEG300, 30% PBS or 30% DI. Scale bars: 50 μm. Asterisks: a rare differentiated cell (A); examples of cells with altered morphology (B). (C) Quantification of the percentage of differentiated cells (% IVL+ cells) of cells grown under the conditions shown in (A, B). (D) Quantification of the percentage of differentiated cells (% IVL+ cells) of cells grown on flat substrates in the presence of different concentrations of PEG 300. Cells were grown for 24h. (C, D) Data are from three experiments in which >1000 cells were analysed per experiment. Experiments were performed with three technical replicates per condition. Statistics: one-way ANOVA (+ Tukey’s) test. Error bars: mean + SD. ** p-value < 0.01. *** p-value < 0.001. (E) Western blot of keratinocytes cultured in suspension for 24h in the presence of 5% PBS (-) or 5% PEG300 (+). Cell lysates were probed with antibodies against involucrin (IVL) to visualise differentiation; antibodies against β-actin were used as a loading control. Full blot and replicates are shown in Supplementary Figure 3.

**Figure 8.**
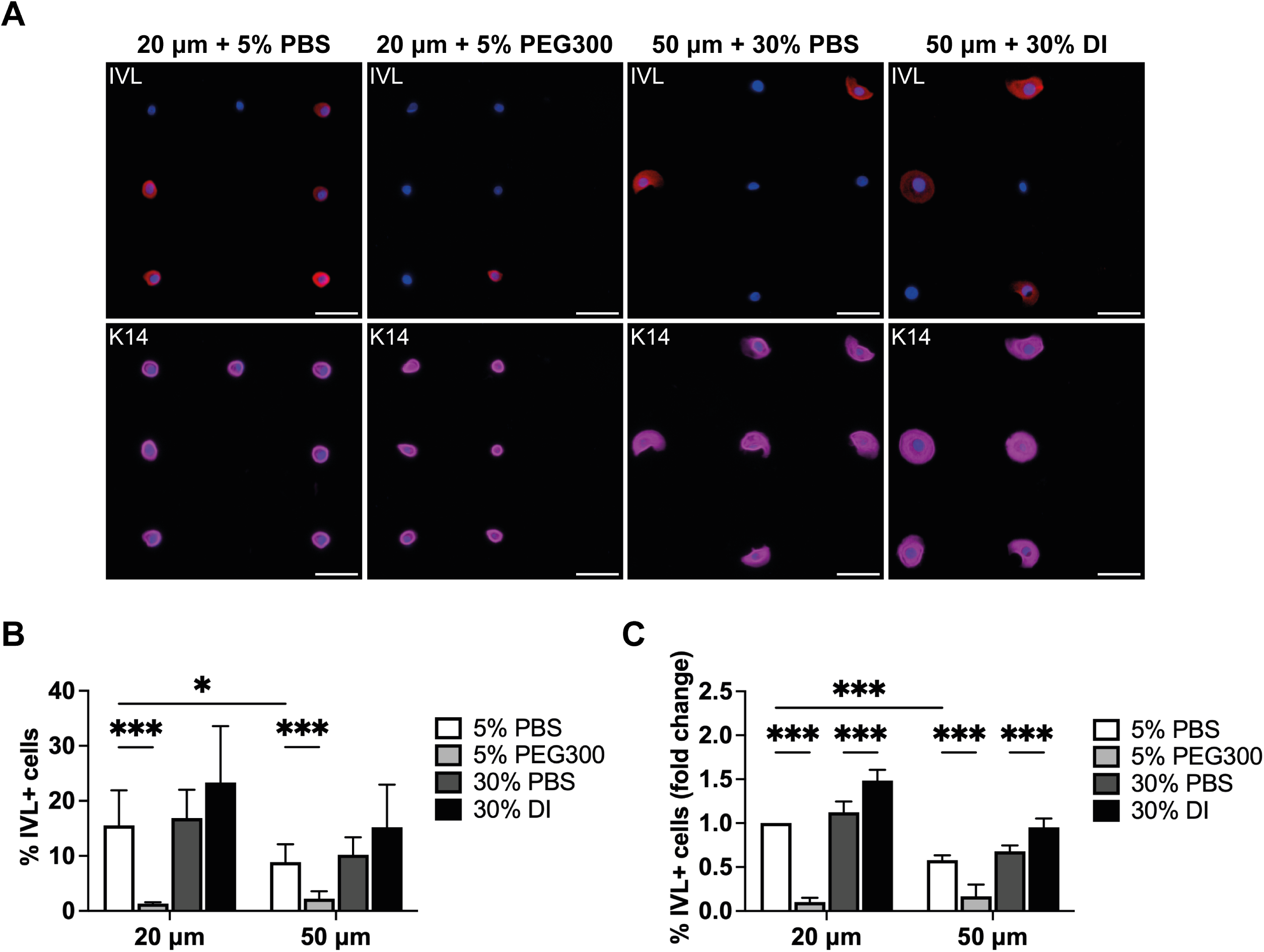
Effects of PEG300 and DI on cells on micropatterned islands. (A) Representative images of anti-involucrin (IVL, red) and anti-keratin14 (K14, magenta) staining of cells grown on 20 μm and 50 μm diameter micropatterned islands. Cells were counterstained with DAPI (blue). Cells were grown in the presence of 5% PBS, 5% PEG300, 30% PBS or 30% DI for 24h. Scale bars: 50 μm. (B-C) Quantification of (B) the percentage of differentiated cells (% IVL+ cells) and (C) the fold change in the percentage of differentiated cells (% IVL+ cells, normalised to 20 μm islands treated with 5% PBS) of cells. Results are from three independent experiments performed with three technical replicates per condition. More than 1000 cells were analysed per experiment. Statistics: two-way ANOVA (+ Šídák’s) test. Error bars: mean + SD. * p-value < 0.05, *** p-value < 0.001.

### Effects of PEG and DI on keratinocytes seeded on S1 and S2 substrates and influence of aquaporin inhibition

We next examined the effects of PEG and DI on cells cultured on S1 and S2 substrates (Figure 9A, B). PEG 300 inhibited differentiation on S1 (Figure 9A), while DI promoted differentiation on S2 (Figure 9 B). On S1, S2 and flat substrates the inhibition of differentiation was reversed by removal of PEG and the resulting % involucrin-positive cells was greater than in the PBS controls (Figure 9A).

**Figure 9.**
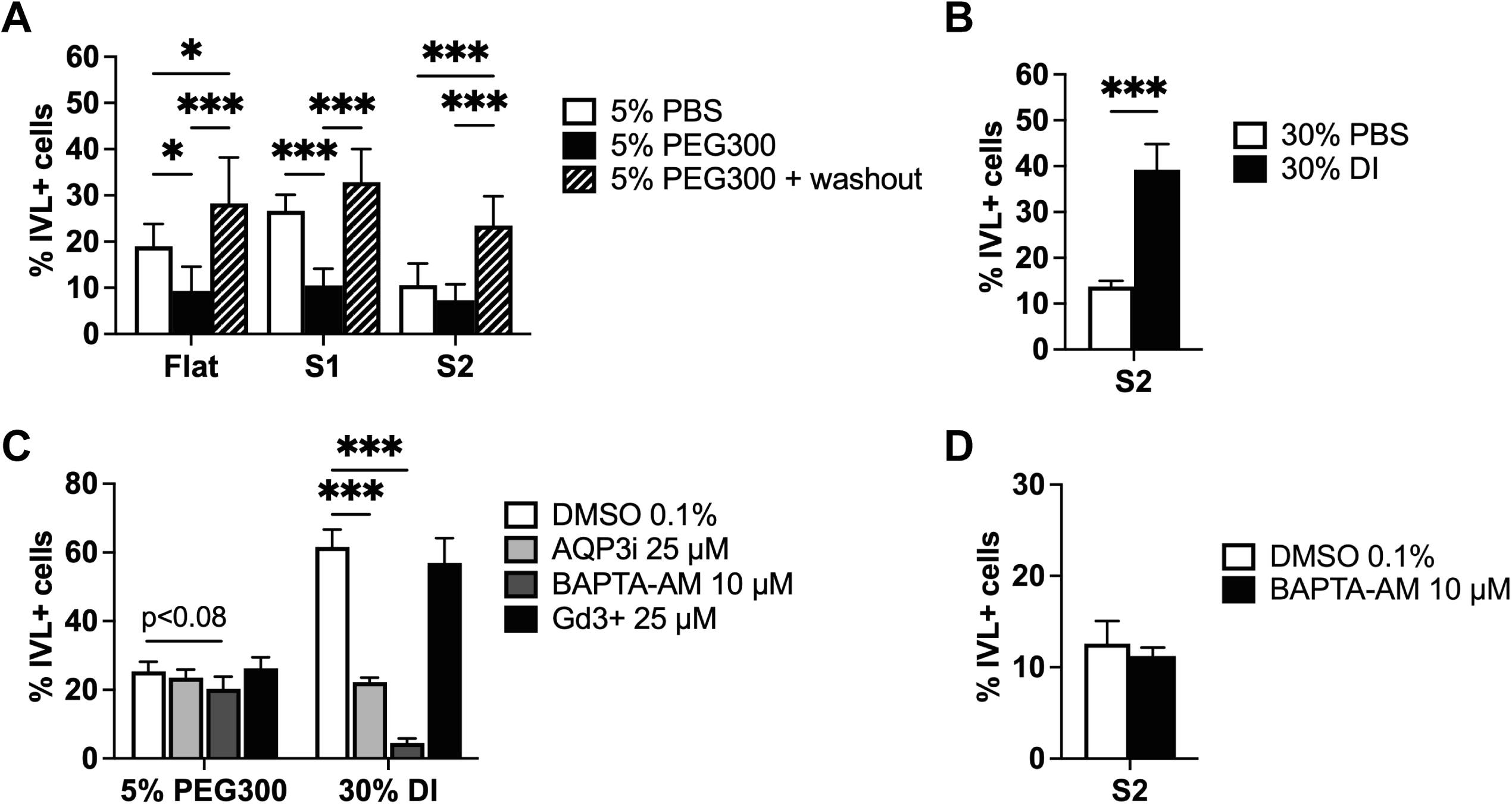
Modulation of differentiation on S1 and S2 substrates. (A) Quantification of the percentage of differentiated cells (% IVL+ cells) on flat, S1 and S2 substrates. Cells were cultured in the presence of 5% PEG 300 for 24h and then fixed or allowed to recover in complete FAD medium for 24h (washout). (B) Cells grown on S2 for 24h in the presence of 30% PBS or 30% DI. Quantification of the percentage of differentiated cells (% IVL+ cells). (C, D) Cells were grown on flat (C) or S2 substrates (D) for 24h in the presence of DMSO (0.1%) or BAPTA-AM (10 μM) (C, D) or 25 μM Gd^3+^ (C) and the percentage of differentiated cells was quantified (% IVL+ cells). Results are from three independent experiments performed with three technical replicates per condition. More than 1000 cells were analysed per experiment in (A, B) and more than 500 cells in (C, D). Statistics: (A, B) one-way ANOVA (+ Tukey’s) test; (C, D) two-tailed unpaired t-test. Error bars: mean + SD. MFI: mean fluorescence intensity. * p-value < 0.05, *** p-value < 0.001.

The plasma membrane of cells is made water permeable through expression of aquaporins (AQPs) and in many cell types swelling is dependent on a rise in intracellular calcium ions (Hoffmann et al., 2009; Jahn et al., 2021). To see which mechanisms could be responsible for the differentiation effects of DI and PEG, cells were treated with a cell permeable analog of 1,2-bis-(2-aminophenoxy)ethane-N,N,N’,N’-tetraacetic acid acetoxymethyl ester (BAPTA-AM) to chelate intracellular calcium (Ca^2+^) (Tsien, 1981; Tang et al., 2007; Nava et al., 2020), with DFP00173 (AQP3i) to inhibit the water transport through aquaporin-3 (Sonntag et al., 2019; de Boer et al., 2023) and with gadolinium (Gd) to block ion channels (predominantly stretch-activated calcium channels and transient receptor protein channels, but also others) (Adding et al., 2001; Bagley et al., 2024). Treatment with BAPTA-AM or AQP3i inhibited the effects of DI on differentiation, whereas Gd did not (Figure 9C, D). These treatments, nonetheless, did not reverse the differentiation inhibitory effects of PEG300 (Figure 9C). BAPTA-AM also did not reverse differentiation on S2 (Figure 9D). Thus, the effects of PEG300 and S2 might be independent of ion channels and intracellular Ca^2+^.

## Discussion

While restricted cell-substrate adhesion is a potent differentiation stimulus for cultured human epidermal stem cells, we have also identified topographies that trigger differentiation of spread cells (Zijl et al., 2019, 2022; Figure 10). In the present report we have compared one topographical substrate that promotes differentiation of spread cells (S1) and one that suppresses differentiation (S2) relative to flat culture substrates. Prompted by the observation that cells on S2 have a reduced volume compared to cells on S1, we examined the effect of modulating keratinocyte volume with PEG 300 and DI. We found that in all contexts tested a reduction in cell volume inhibited differentiation, while an increase stimulated differentiation. Keratinocyte differentiation has long been known to correlate with an increase in cell size (Watt and Green, 1981; Jones and Watt, 1993) but until now there was no evidence for a causative relationship. Our findings lend support to the hypothesis that reduced substrate contact and increased cell volume can synergise to promote keratinocyte differentiation (Watt, 1988).

**Figure 10.**
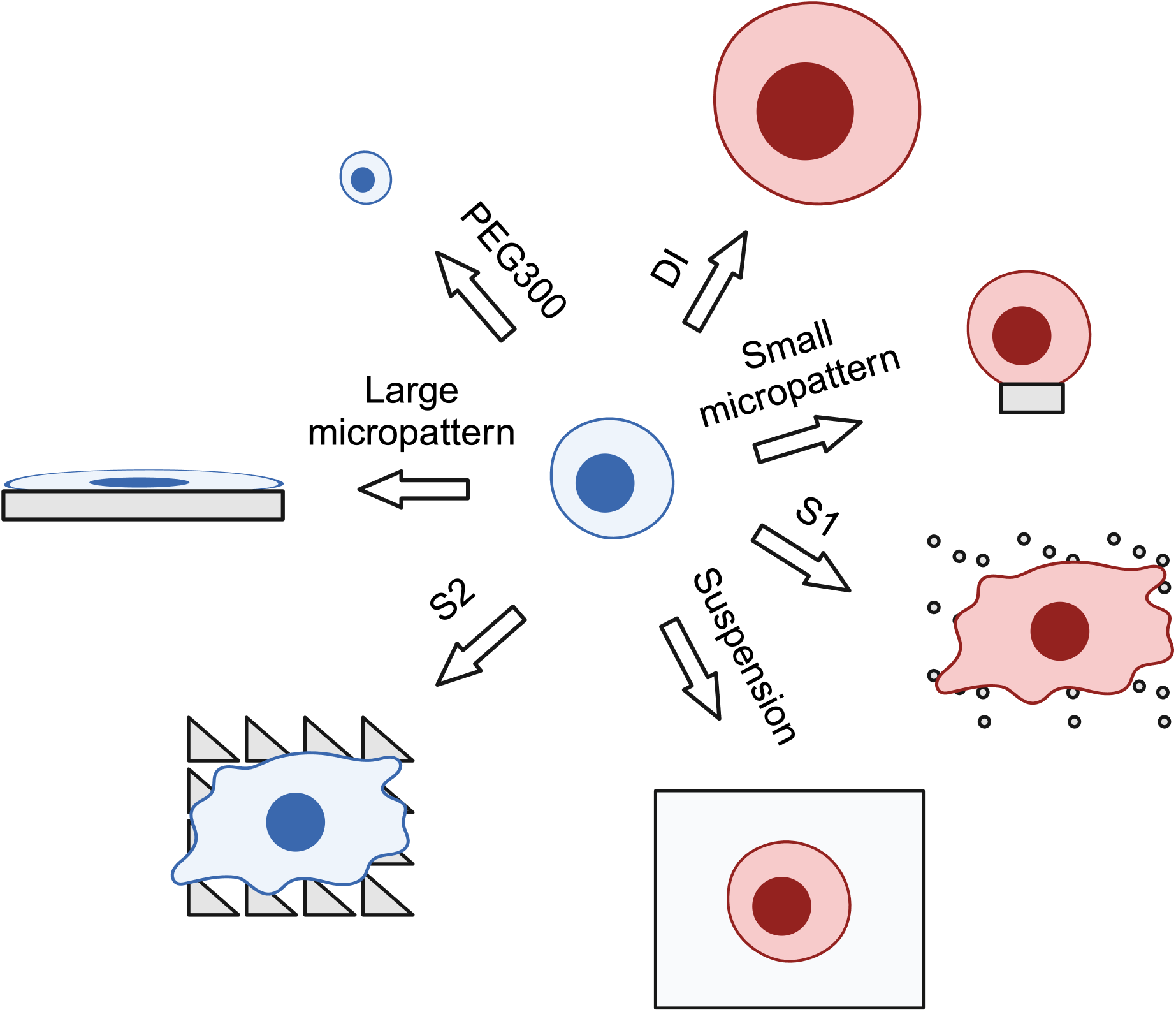
Schematic summary. Cultured epidermal stem cells (blue) can be induced to differentiate (pink) by seeding in suspension, on small micropatterned islands, on S1 substrates or by treatment with DI. The cells remain undifferentiated on S2 substrates, large micropatterned islands or when treated with PEG 300. The effects of PEG 300 and DI are independent of cell-substrate interactions.

Recent studies have demonstrated an association between cell volume and the mechanical properties of the cell cortex and cytoplasm, including cell stiffness (Guo et al., 2017; Jiang and Sun, 2013). Myc-depleted keratinocytes exhibit increased stiffness, which correlates with a decrease in cell area and cortical actin (Bernabé-Rubio et al., 2023) and also reduced epidermal cell size (Zanet et al., 2005). Nevertheless, the correlation between cell stiffness and volume was not observed in our current studies because cell stiffness was lower on both S1 and S2 than on flat substrates, in spite of the reduced cell volume on S2. We have found previously that cell stiffness is affected by the location of keratinocytes on undulating topographies (Mobasseri et al., 2019), leading us to conclude that there are multiple factors that affect keratinocyte stiffness.

When keratinocytes are induced to differentiate in suspension, commitment occurs 4h and upregulation of differentiation markers is evident at 12h (Mishra et al., 2017). Using bulk RNA sequencing we did not detect significant differences in gene expression between keratinocytes plated on S1 and S2 at 1h and 4h, although at 12h genes associated with the differentiation programme were selectively upregulated on S1. Further experiments are necessary to establish whether commitment occurs later on S1 than in suspension and, if so, whether the pro-commitment phosphatase network observed in suspension (Mishra et al., 2017) and in vivo (Reynolds et al., 2021; Negri et al., 2023) is involved. Many transcription factors are mechanosensitive (Bao et al., 2019; Dupont and Wickström, 2022), including SRF/MAL and YAP/TAZ, which are known to regulate keratinocyte differentiation (Zij et al., 2022).

Cells are known to be densely packed with nucleic acids, proteins, lipids and other macromolecules, and this molecular crowding influences many aspects of cell physiology, including cytoskeletal organisation and cell adhesion (Subramanya and Boyd-Shiwarski, 2024). We have previously observed differences in nuclear morphology and cytoskeletal organisation in keratinocytes seeded on S1 and S2 (Zijl et al., 2019) and changes in cell volume are also known to affect molecular crowding (Subramanya and Boyd-Shiwarski, 2024). This leads us to speculate that crowding may play a role in the regulation of epidermal differentiation. While more work needs to be done to explore potential underlying mechanisms it will be interesting to start with the genes that are selectively upregulated on S2 at 12h, including vimentin (Wu et al., 2022) and the microtubule-associated genes ARL4C and CEP170 (Wei et al., 2009; Pillai et al., 2015) (Figure 3).

Cell volume regulation is an integral part of normal physiological processes within tissues (Hoffmann et al., 200). Epidermal cells are able to counteract cell swelling or shrinkage induced by osmotic stress via mechanisms known as regulatory volume decrease or increase (Jahn et al., 2021). Several channels and transporters that regulate keratinocyte volume have been identified (Jahn et al., 2021) and their respective roles in influencing differentiation remain to be examined.

## Methods

### Keratinocyte culture

Primary human keratinocytes (strains km, kn or kp) from neonatal foreskin were cultured on a feeder layer of mitotically inactivated J2 3T3 cells in 1 part Ham’s F12 medium (F12) and 3 parts Dulbecco’s Modified Eagle’s Medium (DMEM) supplemented with 1.8 x 10^-4^M adenine (basal FAD medium). Basal FAD medium was supplemented with 10% foetal calf serum (FCS), 0.5 μg/ml hydrocortisone, 5 μg/ml insulin, 10^−10^M cholera enterotoxin and 10 ng/ml epidermal growth factor to make complete FAD medium. J2-3T3 cells were cultured in DMEM containing 10% donor calf serum.

### Polystyrene topographies

A silicon (Si) wafer template was fabricated using photolithography and etching, as described previously (Zijl et al., 2019; Kelvin Nanotechnology Ltd, UK). The wafer was coated with PDMS (Sylgard 184; Qin et al., 2010) overnight at room temperature and then cured by heating at 80^0^C for at least 5 hours. The cured PDMS was cooled to room temperature and then coated with 25% polystyrene (Goodfellow) dissolved in g-butyrolactone (GBL, Acros Organics; Wang et al., 2011). GBL was evaporated for 4 h at 95 °C, followed by >12 h at 150 °C. To create flat control substrates this same process was performed using flat PDMS. After oxygen plasma treatment (1-2 mins at 0.3-0.4 mBar) (Jokinen et al., 2012) and collagen coating (rat tail collagen type I, 20 μg/mL, overnight at room temperature), the polystyrene surfaces were used for cell seeding. Keratinocytes were suspended in complete FAD medium and seeded at a density of 7.5x 10^4^ cells per cm^2^.

### Scanning electron microscopy

Keratinocytes were grown on collagen-coated polystyrene substrates for 24 hours and chemically fixed in 2.5% (w/v) glutaraldehyde (Science Services, EM grade, E16200) in 0.1M PHEM buffer. Briefly, 2x fixative was added to the cells in a 1:1 ratio with medium and incubated for 15 minutes at room temperature. The 2x fixative was then replaced with fresh 1x fixative, and the cells were incubated for an additional hour at room temperature. The cells were washed twice with 0.1M PHEM buffer and transferred to new wells to remove residual traces of glutaraldehyde. Further fixation was continued with two washes with 0.1M sodium cacodylate buffer. The samples were post-fixed in freshly prepared 1% (w/v) osmium tetroxide (Science Services, E19152), 0.8% potassium-ferrocyanide (Merck, 4984) in 0.1M sodium cacodylate buffer for 45 minutes, protected from light and rinsed four times with dH2O. Afterwards, samples were incubated with 1% (w/v) uranyl acetate (Serva, 77870.01) in dH2O for 30 minutes on ice and subsequently, serially dehydrated in increasing concentration of cold ethanol on a shaker (3x 25%, 2x 50%, 2x 70%, 2x 90%). The samples were then washed twice with 100% ethanol for 10 minutes at room temperature, critical point dried (Leica Microsystems) and mounted on clean SEM stubs with carbon-stickers. The stubs were gold-sputter coated to increase their conductivity (Quorum, QR150 S). Samples were imaged with a Zeiss 540 Crossbeam at 5kV acceleration voltage, a 700pA current, and a secondary electron detector at 0degree (top view) and 54degree tilt (side view) respectively.

### Lentiviral infection

The SV40-puro cassette comprising SV40 promoter and puromycin resistance gene sequences in pLenti-INV3700-mCherry (Hiratsuka et al., 2020) was replaced with the hPGK-LifeAct-EGFP-WPRE sequence from pLenti.PGK.LifeAct-GFP.W (Addgene #51010; Belin et al., 2014) using In-Fusion HD Cloning Kit (Takara). The DNA fragments were obtained by PCR amplification. The sequence of the resulting plasmid (pLenti-INV-mCherry-LifeAct-EGFP) was validated by Sanger sequencing (Source Bioscience Sequencing, Cambridge).

### Flow cytometry

Single cell suspensions of keratinocytes transduced with pLenti-INV-mCherry-LifeAct-EGFP were suspended in phenol-red free complete FAD medium, labelled with DAPI and flow sorted based on the expression of LifeAct-GFP and the absence of IVL-mCherry. Sorting was performed on a FACSAria III Cell Sorter (BD Biosciences). Flow data were analysed using FlowJo (BD Biosciences, version 10). Uninfected cells were used as a negative control to set the GFP+ and mCherry-gates. DAPI was used to exclude dead cells. After sorting, cells were resuspended in pre-warmed complete FAD medium and plated on a J2 feeder layer. When the cultures had reached ∼80% confluence they were harvested for live cell imaging.

### Live cell imaging

pLenti-LifeAct-EGFP-INV-mCherry infected keratinocytes (strain km) were disaggregated and seeded on PS topographies for 45-60 min. Non-adherent cells were removed and adherent cells were incubated at 37 °C for 4-6h before being transferred to phenol red-free FAD medium supplemented with 1 mM sodium pyruvate, 36.5 mM sodium bicarbonate (Gibco) and 100 mM HEPES. Imaging was performed on an upright A1R multi-photon microscope (Nikon) at 37 °C. Cells were imaged at 1h intervals. Images were acquired of multiple z-slices (z-interval: 5-10 μm step size) and combined together in maximum projections. The resolution of the images was 512 x 512 pixels and the magnification was 25x. Cells with an mCherry mean fluorescent intensity (MFI) of >500 were scored as differentiating (active IVL promoter) (Hiratsuka et al., 2020).

### Immunofluorescence staining

To label with antibodies to involucrin (SY3 or SY7 mouse monoclonal antibodies; Hudson et al., 1992) or Keratin 14 (chicken polyclonal antibody; Biolegend 906001/906004) cells were first fixed in 4% (w/v) paraformaldehyde (PFA, Sigma) diluted in phosphate buffered saline (PBS, Sigma) for 15 mins at room temperature then permeabilised in 0.2% (w/v) Triton X-100 (Tx100, Sigma, diluted in PBS for 15-20 mins at room temperature (RT). Samples were washed three times with PBS and blocked in PBS containing 10% (v/v) FBS and 0.25% (w/v) cold water fish skin gelatin (Sigma) (Connelly, 2010) for 1h at room temperature. Primary antibodies were diluted in blocking buffer and incubated with the fixed cells overnight at 4 °C or for 1h at RT. After washing 3x with PBS cells were incubated with Phalloidin-conjugated secondary antibodies in blocking buffer, (Invitrogen A12379) and 4′,6-diamidino-2-phenylindole (DAPI, Invitrogen D1306) for 1h at room temperature in the dark. After staining, samples were washed 3x with PBS and mounted onto glass 6-well plates (Cellvis) with ProLong Gold Antifade Mountant (ThermoFisher) or Mowiol 4-88 (Sigma).

### Confocal microscopy

Immunofluorescence labelled cells on polystyrene substrates were analysed by confocal microscopy. Images at 10x, 20x or 40x magnification were acquired using dry objectives on an A1 upright confocal (Nikon) at the Nikon Imaging Centre (King’s College London). Images at 63x magnification were acquired on an A1R inverted confocal (Nikon), using an oil objective (immersion oil, Nikon).

Confocal images were analysed using ImageJ/Fiji. Mean fluorescence intensity (MFI) values were computed using an automated imaging pipeline and were constructed using Jython (Python implementation in ImageJ/Fiji), as described previously (Louis et al., 2022). Pipelines were used to compute the mean fluorescent intensity values (MFIs) and morphological measurements of individual cells. Subsequent data analysis was done in ImageJ/Fiji and Excel. Unless otherwise stated, intensity values represent mean fluorescent intensity values per cell (MFI). The MFI is the average intensity of all cells in an experiment. To determine the percentage of positive cells for a marker of interest, intensity thresholds were set based on the background intensity (areas in the images without staining for the marker of interest) and cells labelled with a secondary antibody only control. Unless otherwise stated, cells were considered differentiated (IVL+) if their cytoplasmic MFI for IVL was higher than that of cells stained with a secondary antibody control (cells that were not stained with a primary antibody against TGM1/IVL, but only with a secondary antibody). IVL+ cells also had MFIs higher than the background intensity (MFI of areas negative for IVL).

### Cell volume measurements

To quantify the volume of cells grown on polystyrene substrates, cells were fixed with 3.7% formaldehyde or 4% PFA in PBS, labelled with anti-keratin 14 to mark the cytoplasm and DAPI to mark nuclei, and imaged by confocal microscopy. The z-slice interval was set at 0.5 μm (Hansen et al., 2022; Koushki et al., 2023). Confocal measurements of cell volume have been shown to be similar to super resolution imaging measurements (Guo et al., 2017; Lee et al., 2019, 2021). Z-stacks were deconvoluted using the NIS elements imaging software (Nikon), and cell and nuclear volumes were calculated in ImageJ/Fiji (NIH, voxel counter plug-in) (Koushki et al., 2023). Thresholding (default method) was performed to exclude background staining. Total thresholded volumes were divided by the number of cells per image, to obtain the average cell/nuclear volume per cell for each image. Averages for different images were combined together to create averages for different experiments.

### Cell volume manipulation

Cells were treated with 5% (v/v) PEG300 or 30% (v/v) DI (Cai et al., 2019; Guo et al., 2017; Li et al., 2021; Rashid et al., 2023; Tomba et al., 2022). Cells treated with 5% (v/v) PBS (for 5% PEG300) and 30% (v/v) PBS (for 30% DI) were used as controls. For cell volume manipulation on PS substrates, cells were allowed to attach for 45-60 mins and non-adherent cells were removed. The medium was switched to FAD medium containing 5% PBS, 5% PEG300, 30% PBS or 30% DI. Cells were cultured for a further 24h to allow differentiation to occur (Connelly, 2010) and then fixed and stained for analysis.

For cell volume manipulation on unpatterned substrates, cells were seeded on collagen-I coated (20 μg/mL in PBS) 96-well plates (Greiner, μClear black) at a density of 1.0-1.5 x 10^4^ cells/well (3.0-5.0 x 10^4^ cells/cm^2^) for 45-60min. After removal of non-adherent cells, the cells were transferred to complete FAD containing 5% PBS, 5% PEG300, 30% PBS or 30% DI and cultured for 24h. before fixation and labelling.

### Pharmacological inhibitors

To chelate intracellular calcium, cells were treated with a cell permeable analog of 1,2-bis-(2-aminophenoxy)ethane-N,N,N’,N’-tetraacetic acid acetoxymethyl ester (BAPTA-AM, Abcam, 10 μM) (Tsien, 1981; Tang et al., 2007; Nava et al., 2020). Gadolinium trichloride (Gd^3+^, Sigma, 25 μM) was used to block ion channels (predominantly stretch-activated calcium channels and transient receptor protein channels, but also others) (Adding et al., 2001; Bagley et al., 2024). DFP00173 (AQP3i, Cambridge Bioscience, 25 μM) was used to inhibit the water transport through aquaporin-3 (Sonntag et al., 2019; de Boer et al., 2023). All drugs were diluted in dimethyl sulfoxide (DMSO, Sigma). Cells treated with DMSO only (vehicle) were used as negative controls (maximum final concentration of DMSO: 0.1%).

### Differentiation on micro-patterned islands and in suspension

20 μm and 50 μm diameter micropatterned islands were prepared using custom Quartz photomasks (JD Photodata) as described previously (Louis et al., 2022). To induce terminal differentiation in suspension, pre-confluent keratinocytes were disaggregated and resuspended at a concentration of 10^5^ cells per ml in complete FAD medium supplemented with 1.45% methylcellulose (4,000 centipoises, Aldrich). The cell suspensions were then plated in 35-mm-diameter bacteriological plastic dishes coated with 0.4% polyHEMA. After incubation for up to 24 h at 37 °C the methylcellulose was diluted with PBS and the cells recovered by centrifugation (Louis et al., 2022).

### RNA sequencing and RT-qPCR

RNA isolation was performed using the RNeasy mini kit (Qiagen), according to the manufacturer’s instructions. For real-time quantitative PCR (RT-qPCR), complementary DNA (cDNA) was synthesized using the Quantitect Reverse Transcription kit (Qiagen), according to the manufacturer’s instructions. The cDNA was mixed with Fast SYBR Green (SYBR, Applied Biosystems), nuclease-free water, and custom-made oligonucleotide primers (Sigma). Reactions were performed in 384-well PCR plates (Bio-Rad) using a CFX384 Touch Real-Time PCR Detection System (Bio-Rad). The primers used for Q-PCR are listed in Supplementary Table 2. Each sample was run with four technical replicates and three experimental replicates per condition. Gene expression was quantified using the 2-ΔΔCT method (Livak and Schmittgen, 2001). Cycle threshold (Ct) values were averaged across the different technical replicates. Ct values were normalised to the expression of housekeeping genes (the average of 18S, GAPDH and TBP). Gene expression in different samples was normalised to control conditions (flat substrates). Melting curves for the different primers were checked after each experiment. Primers that generated unusual melting curves (e.g. melting curves with several peaks) were replaced. Primers that generated Ct values in negative control reactions (nuclease-free H2O instead of cDNA) were also replaced.

To prepare keratinocytes for bulk RNA sequencing (RNA seq), RNA was isolated using the RNeasy mini kit (Qiagen). Each experiment was performed with three technical replicates per condition. Technical replicates were pooled together. The quality of the RNA was checked on a Qubit 2.0 Fluorometer (Thermo Fisher). Samples submitted for sequencing had an RNA integrity number of 9.9-10.0. Messenger RNA (mRNA) selection, library preparation and sequencing were performed by GENEWIZ Inc., as described (Cujba et al., 2022) according to standardized Illumina protocols. High throughput RNA seq was performed on a NovaSeq 6000 Sequencing System (Illumina). At least 2.0 x 10^7^ paired-end reads of 150 bp were obtained per sample. The mean quality score of the samples was >35, as determined by FastQC.

Raw RNA sequencing files were processed by GENEWIZ. To remove possible adapter sequences and poor quality nucleotides, sequencing reads were trimmed with Trimmomatic v.0.36 (Bolger et al., 2014). Quality control was performed using FastQC/0.11.8 (https://www.bioinformatics.babraham.ac.uk/projects/fastqc/). Trimmed reads were mapped to the human GRCh38 reference genome, using STAR aligner v.2.5.2b (Dobin and Gingeras, 2015). Transcript abundance was calculated using ‘featureCounts’ from the ‘Subread’ package v.1.5.2 (Liao et al., 2013). Only unique reads that fell within exon regions were counted. Gene expression analysis was performed in R version 3.5.5, using ’DESeq2’ (Love et al., 2014). The Wald test was used to generate p-values and log2 fold changes (LFCs). Genes with a Benjamini and Hochberg adjusted p-value (padj) < 0.05 and an absolute log2 fold change > 1 (or < -1) were considered significantly differentially expressed. Volcano plots were generated using the ‘EnhancedVolcano’ package. Heatmaps were made using the heatmap.2 function from the ‘gplots’ package and plotted by me in GraphPad Prism. Principal component analysis (PCA) was performed using the plotPCA function in DESeq2.

### Atomic force microscopy

Keratinocytes were cultured on PS substrates for 12h and then transferred to FAD medium containing 100 mM HEPES. Measurements were performed on a BioScope Resolve BioAFM (Bruker), which was coupled to an optical microscope (Nikon Eclipse Ti-U). Measurements were carried out on live cells at 37 ^0^C using a spherical nitride tip (5 μm) and nitride cantilever (SAA-SPH-5UM, Bruker). For each sample, 24×24 force extension measurements were performed to probe stiffness. Each measurement involved a 10 μm piezo-electric excursion up to a maximum force of 10 nN. The Young’s modulus (cell stiffness) was calculated by fitting the force-extension curves with a Hertzian model (spherical), assuming a Poisson’s ratio of 0.5, using the Nanoscope software (version 1.8, Bruker). Only the region between 30% and 70% of the maximum force was employed for fitting. For each experiment, 10 randomly selected cells were imaged per condition (flat, S1 and S2). Twenty force curves were generated per cell. Experiments were repeated three times.

### Western blotting

Cells were lysed for 30 min on ice in RIPA buffer (Sigma) containing Pierce Protease Inhibitor (Thermo Scientific), then sonicated for 3 x 25s at 4 °C (CamSonix C080T Ultrasonic bath, Camlab). Lysates were centrifuged at 16,000 rcf for 10 min at 4 °C and the pellets discarded. The protein concentrations of supernatants were determined using a Pierce BCA Protein Assay Kit (Thermo Scientific), according to the manufacturer’s instructions.

For western blotting, 25 μg of protein was diluted in Laemmli SDS sample buffer (Thermo Scientific), boiled at 95^0^C for 5 min and loaded into each lane of a 4–15% Mini-PROTEAN TGX Stain-Free precast gel (Bio-Rad). Colour Prestained Protein ladder was loaded in some wells in order to determine the molecular mass of proteins in the experimental samples. Gels were submerged in 1x Tris/glycine/SDS running buffer (Bio-Rad) and subjected to SDSPAGE for approximately 2h at 100-120V. Afterwards, gels were transferred to PVDF membranes in Trans-Blot Turbo Mini 0.2 μm PVDF Transfer Packs (Bio-Rad) using the Trans-Blot Turbo Transfer System (Bio-Rad, 7 mins at 20V).

Membranes were blocked for 1h at room temperature in Tris-balanced saline containing 0.025% Tween-20 (0.025% TBST, Severn Biotech) and 5% non-fat skimmed milk (Tesco) then incubated with primary antibodies diluted in blocking buffer overnight at 4^0^C. After washing 3x with 0.025% TBST for 10 min membranes were incubated with secondary antibodies conjugated to horseradish peroxidase in blocking buffer overnight at 4^0^C. Following 3 x 10 min washes in 0.025% TBST and blots were developed using Clarity Western ECL Substrate (Bio-Rad), according to the manufacturer’s instructions. Protein bands were visualised on a ChemiDoc Touch Imaging System (Bio-Rad). The following antibodies were used: SY7 mouse anti-involucrin (Hudson et al., 1992), HRP conjugated mouse anti-β-actin (Santa Cruz sc-47778 HRP) and HRP conjugated horse anti-mouse IgG (Cell Signaling Technology 7076).

### Statistics and reproducibility

Graphing was performed in Excel or Graphpad Prism (Dotmatics). Statistical analyses were performed in GraphPad Prism. Multiple testing corrections were used where indicated - e.g. one-way ANOVA [+ Tukey’s] test means a Tukey’s multiple comparisons test was used in combination with a one-way ANOVA test. Only significant (p-value < 0.05) and near significant p-values (p-value < 0.10) are shown. Error bars represent the standard deviation (SD) from the mean (mean + SD) unless otherwise stated. Images are representative of the indicated number of experiments. Generally, experiments were performed with three experimental replicates per condition per substrate and three technical replicates per condition (e.g. per time point), unless otherwise stated.

## Acknowledgements

This work was funded by grants to FMW from the UK Medical Research Council (MR/PO18823/1), the Wellcome Trust (206439/Z/17/Z; 098503/Z/12/Z) and the Danish National Research Foundation (DNRF135). FMW gratefully acknowledges financial support from EMBO and technical assistance from the EMBL Electron Microscopy Core Facility, the King’s College London Nikon Imaging Centre and the King’s College London Flow Cytometry Core Facility. SGM was supported in part by the Francis Crick Institute which receives its core funding from Cancer Research UK (CC0102), the UK Medical Research Council (CC0102) and the Wellcome Trust (CC0102). This work was also supported by a EPSRC Strategic Equipment Grant (EP/M022536/1), BBSRC sLoLa (BB/V003518/1), Leverhulme Trust Research Leadership Award (RL 2016-015), Wellcome Trust Investigator Award (212218/Z/18/Z) and Royal Society Wolfson Fellowship (RSWF/R3/183006) to SGM.

## Supplementary Figures and Tables

**Supplementary Figure 1.** Gene ontology (GO) analysis of differentially expressed genes expressed between S1 and S2 at 12h (LFC > 1 or LFC < -1, padj <0.05). Differentially expressed genes were clustered and tested for GO enrichment using GeneSCF v1.1-p2 (Subhash & Kanduri, 2016). GO term enrichment was statistically tested (Fisher’s exact test) and significant p-values were adjusted for multiple testing (p-adjusted). The top 40 most significantly enriched GO terms (p-adjusted < 0.05) are shown. GO terms (y-axis) are plotted against the -log10 of their adjusted p-value (p-adjusted).

**Supplementary Figure 2.** Real-time quantitative PCR of differentially expressed genes. Quantification of messenger RNA (RT-qPCR) expression of genes identified by bulk RNA sequencing as being differentially upregulated on S1 (A) or S2 (B) substrates. Cells were cultured on flat, S1 or S2 substrates for 12h. Statistics: Kruskal-Wallis (+ Dunn’s test). RFE: relative fold expression (compared to flat substrates). * p-value < 0.05, ** p-value < 0.01, *** p-value < 0.001.

**Supplementary Figure 3.** Western blotting of cells placed in suspension for 24h to induce terminal differentiation. The starting cell populations served as controls (Ctrl). Cells were suspended in the presence or absence of 5% PEG 300, then lysed and subjected to western blotting for involucrin (IVL), β-actin or GAPDH (loading controls). Each blot includes colour prestained protein ladders (two lanes on left hand side and two lanes on right hand side; Bio-Rad). Lysates from four independent experiments (Exp.) are shown. Red boxes show regions included in Figure 7E.

**Supplementary Table 1.** Differential gene expression of cells on S1 and S2 at 12h. The table shows significantly differentially expressed (LFC > 0.5 or < -0.5, padj < 0.05) genes. ENSEMBL gene identification number (ID) and names are shown. Green shading denotes genes upregulated on S2; red shading denotes genes downregulated on S2.

**Supplementary Table 2.** Oligonucleotide primers used in RT-qPCR reactions.

